# Dynamic response of RNA editing to temperature in Grape by RNA deep-sequencing

**DOI:** 10.1101/745364

**Authors:** Aidi Zhang, Xiaohan Jiang, Fuping Zhang, Tengfei Wang, Xiujun Zhang

## Abstract

RNA editing is a post-transcriptional process of modifying genetic information on RNA molecules, which provides cells an additional level of gene expression regulation. Unlike mammals, in land plants, RNA editing converts C to U residues in organelles. However, its potential role in response to different stressors (heat, salt and so on) remains unclear. Grape is one of the most popular and economically important fruits in the world, and its production, like other crops, must deal with abiotic and biotic stresses, which cause reductions in yield and fruit quality. In our study, we tested the influence of the environmental factor temperature on RNA processing in the whole mRNA from grape organelle. In total, we identified 123 and 628 RNA editing in chloroplast and mitochondria respectively with the average editing extent nearly ~60%. The analyses revealed that number of non-synonymous editing sites were higher than that of synonymous editing sites, and the amino acid substitution type tend to be hydrophobic. Additionally, the overall editing level decreased with the temperature rises, especially several gene transcripts in chloroplast and mitochondria (*matK, ndhB* etc.). 245 sites were furthermore determined as stress-responsive sites candidates. We also found that the expression level of PPR genes decreased with the temperature rises, which may contribute to the loss of RNA editing at high temperature. Our findings suggest that the RNA editing events are very sensitive to high temperature, the changes of amino acid in these genes may contribute to the stress adaption for grape.

## Introduction

RNA editing is a post-transcriptional process that could alter the nucleotide sequence of gene transcripts, potentially diversifies the transcriptome and proteomes beyond the genomic blueprint (Takenaka et al. 2013). In mammals, RNA editing occurs in nucleus transcripts, adenosine-to-inosine (A-to-I) editing is the most abundant type of RNA editing catalyzed by the protein family called adenosine deaminases acting on RNA (*ADAR*). Mutants lacking the *ADAR* enzyme exhibit behavior alterations including defects in flight, motor control, and mating (Palladino et al. 2000). However, in plant, RNA editing primarily occurs in organelle transcripts, and the RNA editing type is dominated by cytosine-to-uracil (C-to-U) RNA editing; in ferns and mosses, it also changes U nucleotides to C nucleotides. RNA editing was first documented over a decade ago in mitochondria as sequence differences between DNA and RNA (Covello and Gray 1989; Gualberto et al. 1989; Hiesel et al. 1989), then a number of editing sites in both two organelles (plastids and mitochondria) were subsequently reported in all land plants, including all major plant lineages from the bryophytes to gymnosperms and in all angiosperms. Numerous studies have demonstrated that pentatrico peptide repeat (PPR) proteins play a central role in the process of RNA editing by interacting with several additional nonPPR protein factors, such as multiple organelle RNA editing factors (MORF) (Hammani and Giege 2014), termed as editosome machinery. PPR proteins have been reported to contain conserved domains to edit mRNA of organelle genes and have formed an extended family during plant evolution.

RNA editing has various biological functions(Fujii and Small 2011; Hammani and Giege 2014), recent studies reported that RNA editing plays important roles in various plant developmental processes and evolutionary adaptation, including organelle biogenesis, signal transduction and adaptation to environmental changes. Accordingly, numerous evidences have reported that RNA editing on transcripts of chloroplast and mitochondrial to be responsive to various environmental stressors, especially temperature, salt, affected editing of select transcripts, and the RNA second structure may regulate this relationship (Garrett and Rosenthal 2012a; Garrett and Rosenthal 2012b; Karcher and Bock 2002; Kurihara-Yonemoto and Handa 2001; Kurihara-Yonemoto and Kubo 2010; Rieder et al. 2015; Riemondy et al. 2018; Rodrigues et al. 2017). It was demonstrated that many mutants with impaired editing of specific sites exhibited strong deleterious phenotypes, even lethality (Tang et al. 2017). Additionally, many *PPR* gene mutants can further alter their morphological appearances under stress conditions compared to the wild-type (Tan et al. 2014; Tang et al. 2017). Therefore, RNA editing is likely playing a role in plant resistance to a given environmental stress, then it is necessary to explore how this influences plant responses to environmental stresses. Since the advent of next generation sequencing technologies, RNA-seq becomes a comprehensive, precise, and low-cost approach for transcriptome profiling and variant analysis. Then ten of thousands editing sites have been identified in more and more plants. The growing of public RNA-seq data also provides an excellent opportunity to investigate the effect of RNA editing on organelle function and evolution (Smith 2013). Hundreds of the editing sites are located in coding sequences, and many of them alter protein sequence, thus expanding the proteome diversity and increasing genomic flexibility.

Grape (*Vitis vinifera L.*) is one of the most popular and economically important fruits in the world. Grape production, like other crops, however, must deal with abiotic and biotic stresses, which cause reductions in yield and fruit quality. It remains unclear that whether the RNA editing response to various stresses, such as temperature, water and so on. In this study, we tested the influence of the environmental factor temperature on RNA processing based on whole mRNA deep-sequencing data. Hundreds of RNA editing sites were identified in chloroplast and mitochondria respectively with the average editing extent nearly ~60%. Additionally, we also found a series of characteristics for the editing sites, such as the number of non-synonymous editing sites were higher than that of synonymous editing sites, and the amino acid substitution type tend to be hydrophobic. Notably, the overall editing level decreased with the temperature rises, especially several gene transcripts in chloroplast and mitochondria (*matK, ndhB, nad, atp etc.*). In addition, we found that the expression of the editing enzyme, PPR proteins is dramatically decreased at elevated temperatures, partially, but not fully, explaining the RNA editing responses to temperature. Our results suggest that the RNA editing process responding to temperature alterations maybe due to temperature sensitive expression or stability of the RNA editing enzyme. Environmental cues, in this case temperature, rapidly reprogram the grape transcriptome through RNA editing, presumably resulting in altered proteomic ratios of edited and unedited proteins. Our findings also suggest that the changes of amino acid in these genes may contribute to the stress adaption for grape.

## Material and Methods

### Data collection

All data sets used in this study are publicly available. The RNA-Seq data of grape leaves that treated with different temperatures (25°C, 35°C, 40°C, and 45°C) were downloaded from https://www.ncbi.nlm.nih.gov/bioproject/PRJNA350310. The genome sequence of grape mitochondria and chloroplast and corresponding genome annotation files in ‘tbl’ format were downloaded from the NCBI data repository (https://www.ncbi.nlm.nih.gov; accession numbers: NC_012119.1, NC_007957.1). The whole genome and annotation file of grape was also downloaded from the NCBI data repository for expression analysis (Jaillon et al. 2007).

### Pre-analysis of transcriptome data

We performed the RNA editing sites identification for grape mitochondria and chloroplast separately, and the identification process was split into two steps. For each genome, firstly, we aligned the transcriptome data against the genome, secondly, the RNA editing sites were identified based on the called SNPs-calling results. In order to increase the sequencing depth, we merged the three duplicates under one condition into one sample. Take the mitochondria genome as an example, the quality control of paired-end Illumina sequencing data were firstly evaluated by NGSQCToolkit and low-quality sequence data were filtered out (cutOffQualScore<20) (Patel and Jain 2012), and the treated cleaned reads were aligned to the reference mitochondria genome using the HISAT2 software (default parameters) (Kim et al. 2015). Samtools (Li et al. 2009) was used to index, merge, sort, remove, format convert, mpileup and remove duplications against the aligned data. Afterwards, the bcftools was used to perform SNP-calling based on the treated bam files, and the VCF files were generated (Narasimhan et al. 2016) for Subsequent analysis.

### Identification of RNA editing sites

Based on the SNP-calling results (in ‘VCF’ format) and genome annotation files (in ‘tbl’ format). We utilized the REDO tool to identify the RNA editing sites under default parameter values (Wu et al. 2018). REDO is a comprehensive application tool for identifying RNA editing events in plant organelles based on variant call format files from RNA-sequencing data. REDO only uses the variant call format (VCF) files (records for all sites), the genome sequence file (FASTA format), and the gene annotation file (feature table file in ‘tbl’ format, www.ncbi.nlm.nih.gov/projects/Sequin/table.html) as input. Then the raw variants are filtered by ten rule-dependent filters and statistical filters to reduce the false positives as the following steps, (1) quality control filter, (2) depth filter (DP>4), (3) alt proportion filter (alt proportion <0.1), (4) multiple alt filter, (5) distance filter, (6) spliced junction filter, (7) indel filter, likelihood ratio (LLR) test filter (LLR <10), (8) fisher exact test filter (p value<0.01), (9) complicated filter model. Finally, the RNA editing sites and corresponding annotation information files were generated.

### Characteristic statistics and identified RNA editing sites

For each organelle, the resulted editing sites are used for further statistics and feature analysis, including statistics of editing number, editing type, codon position, amino acid changes, involved genes and so on. Furthermore, we compared the RNA editing efficiency between each two conditions, and identified the editing sites with statistical significance (p value < 0.01). A clustering analysis and heatmap plotting were also performed for each organelle to decipher the tendency of RNA editing extent based on the RNA editing proportion. Genes with significantly changes in editing sites were picked out for further functional analysis.

### Expression analyses of PPR genes

Transcriptome analysis of RNA-Seq data used in our study was also performed for measuring and comparing the levels of gene expression of PPR genes. The protocol that described in previous study (Pertea et al. 2016) was used. Concretely, treated reads from each sample were mapped to the reference genome with HISAT2 (default parameters), stringtie was used for transcript assembly, samtools was used to index, merge, sort, remove, format convert, mpileup and remove duplications against the aligned data; finally, ballgown was used to determines which genes and transcripts are deferentially expressed between each two conditions.

## Results

### Alignment of transcriptome data

There were a total of 12 RNA-Seq samples that treated with different temperatures (25°C, 35°C, 40°C, and 45°C) in our study, each condition has three replicates. We aligned the transcriptome data to the organelle reference genome respectively. The size of mitochondrion genome is about 773,279 bp, encoding 158 genes, for each sample, there were about 7,000 reads mapped to reference with mapping rate about 0.45% (std = 0.185). For chloroplast, the size of its genome is about 160,928 bp, encoding 120 genes. there were about 3,000 reads mapped to reference with mapping rate about 2.13% (std=0.7681),. The higher mapping rate of chloroplast maybe due to its smaller genome size. The statics of reference-guided mapping rate was shown in Fig.1 and Table 1.

**Fig.1.**
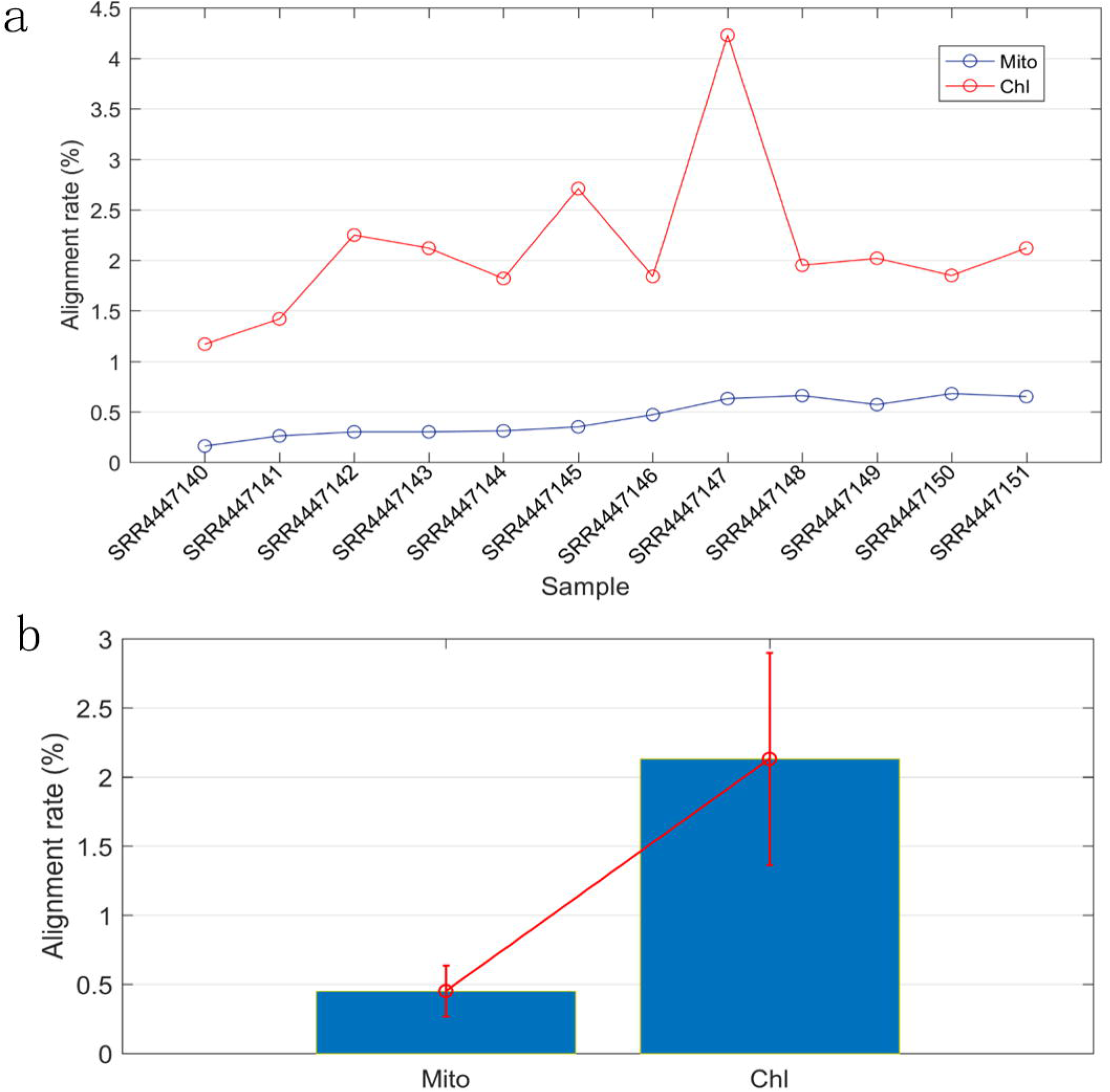
The mapping rates of transcriptome data to mitochondrion and chloroplast genomes. a The statics of reference-guided mapping rate. The y axis represents mapping rate of each sample, and the x axis represents each sample. b The bar figure of The statics of reference-guided mapping rate. The y axis represents mapping rate of each sample, and the x axis represents organelle (Chl: chloroplast; and Mito: mitochondria).

**Table 1.**
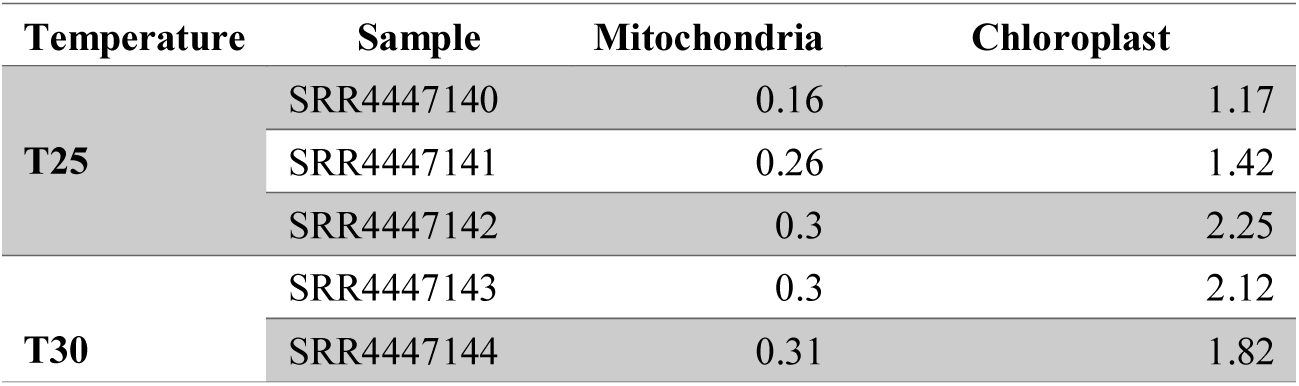
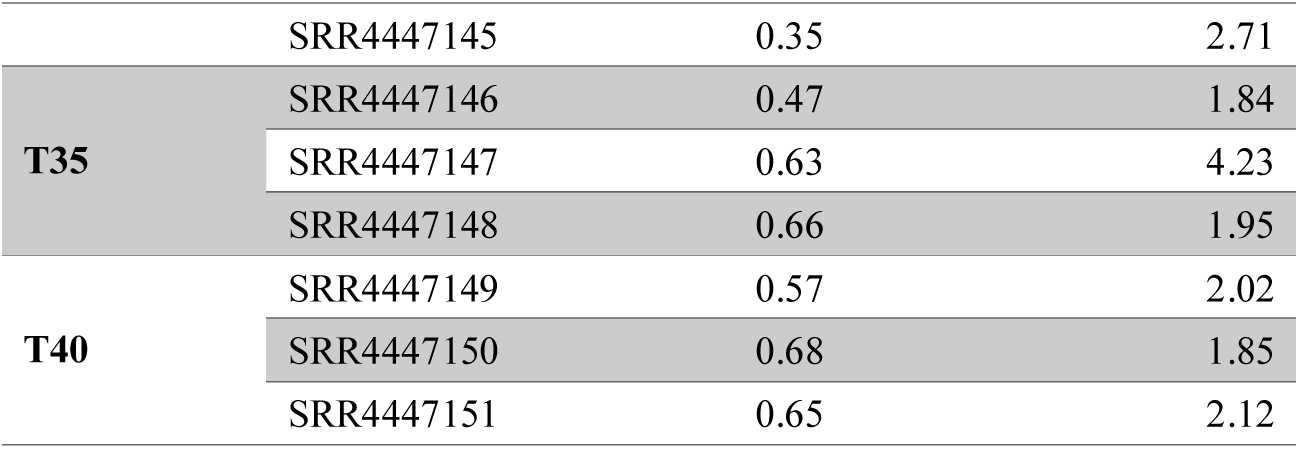
Summary of mapping rates of transcriptome data

### Identification of RNA editing sites

In order to increase the sequencing depth and reliability of editing sites, the resulting bam files of three replicates under one condition were merged for subsequent identification of RNA editing sites. Based on the SNP-calling results and organelle genome annotation file, a total of 751 RNA editing sites were identified in both organelles, however, a few editing sites only appeared under certain condition owning to bias sequencing depth and strict quality control. Take samples at 25°C temperatures in chloroplast as illustration, the attributes of RNA editing sites was shown in Fig. S1. For mitochondrion, there were 627 RNA editing sites identified, involving 53 gene, the number of RNA editing sites identified at different temperatures (25°C, 35°C, 40°C, and 45°C) correspond to 468, 509, 563, 582, along with the increment of temperature, the number of sites increased obviously, as shown in Table 2. In contrast, there were only 122 editing sites identified in chloroplast, and the number of sites didn’t appear to be rising along with temperature increment, 95 editing sites were identified under three conditions (25°C, 35°C, 40°C), only 82 sites were identified under 45°C temperature. As so far, there are no heat-stress response of RNA editing in mitochondrion, the response for heat stress in mitochondrion observed in our study may affect heat-related genes expression through the way of RNA editing. The statistics results showed that most of editing sites were C-to-U, for chloroplast, 97 out of 122 editing sites was C-to-U, the second-most type was G-to-A (5 out of 122); for mitochondrion, 602 out of 627 editing sites was C-to-U, the second-most type was A-to-G (5 out of 122). The detailed information of RNA editing sites in all chloroplast and mitochondrion samples were shown in Table S1 and Table S4.

**Table 2.** The statistics of identified RNA editing sites in mitochondrion and chloroplast

| **Organelle** | **Samples** | **Total** | **Phase(1,2,3)** | **Silent** | **NonSilent** |
| --- | --- | --- | --- | --- | --- |
| **Chl** | **T25** | 95 | 14,77,12 | 6 | 89 |
|  | **T35** | 95 | 11,76,16 | 9 | 86 |
|  | **T40** | 95 | 15,74,36 | 10 | 85 |
|  | **T45** | 82 | 14,66,31 | 8 | 74 |
| **Mito** | **T25** | 468 | 137,291,62 | 49 | 419 |
|  | **T35** | 509 | 156,305,73 | 59 | 450 |
|  | **T40** | 563 | 170,342,84 | 63 | 500 |
|  | **T45** | 582 | 175,355,84 | 61 | 521 |

### Characteristics of the statistics for RNA editing sites

Additionally, we found that RNA editing occurred in second codon position was mainly the largest in both organelles, followed by first codon position except three samples (35°C, 40°C, and 45°C) of chloroplast, as shown in Fig. 2. In mitochondrion, globally, 30, 58, and 12% of the 627 identified editing sites were found at first, second, and third codon positions, respectively, in agreement with previous study (Edera et al. 2018). Similarly, in chloroplast, 14, 76, and 10% of the 122 identified editing sites were found at first, second, and third codon positions, respectively. Furthermore, the statistics of editing type showed that The majority (~ 95%) of the editing events resulted in non-synonymous codon changes, interestingly, we found that the amino acids changes tend to be hydrophobic, the change from hydrophilic to hydrophobic was the highest, followed by the change from hydrophobic to hydrophobic, take T25 for an example, the proportion of hydrophobic2hydrophobic:hydrophilic2hydrophilic,hydrophobic2hydrophilic,hydrophilic2hydrophobic was 114:49:36:206 in mitochondrion, and 13:9:2:62 in chloroplast. In addition, about ~ 55% of the amino acid changes were hydrophilic2hydrophobic produced by editing sites mainly at second codon positions. The most amino acid changes were Ser-to-Leu and Pro-to-Leu, Serine is hydrophilic, whereas Leucine and Proline are both hydrophobic. These above results are in good agreement with previous studies, which demonstrated that the RNA editing causes an overall increase in hydrophobicity of the resulting proteins.

**Fig.2.**
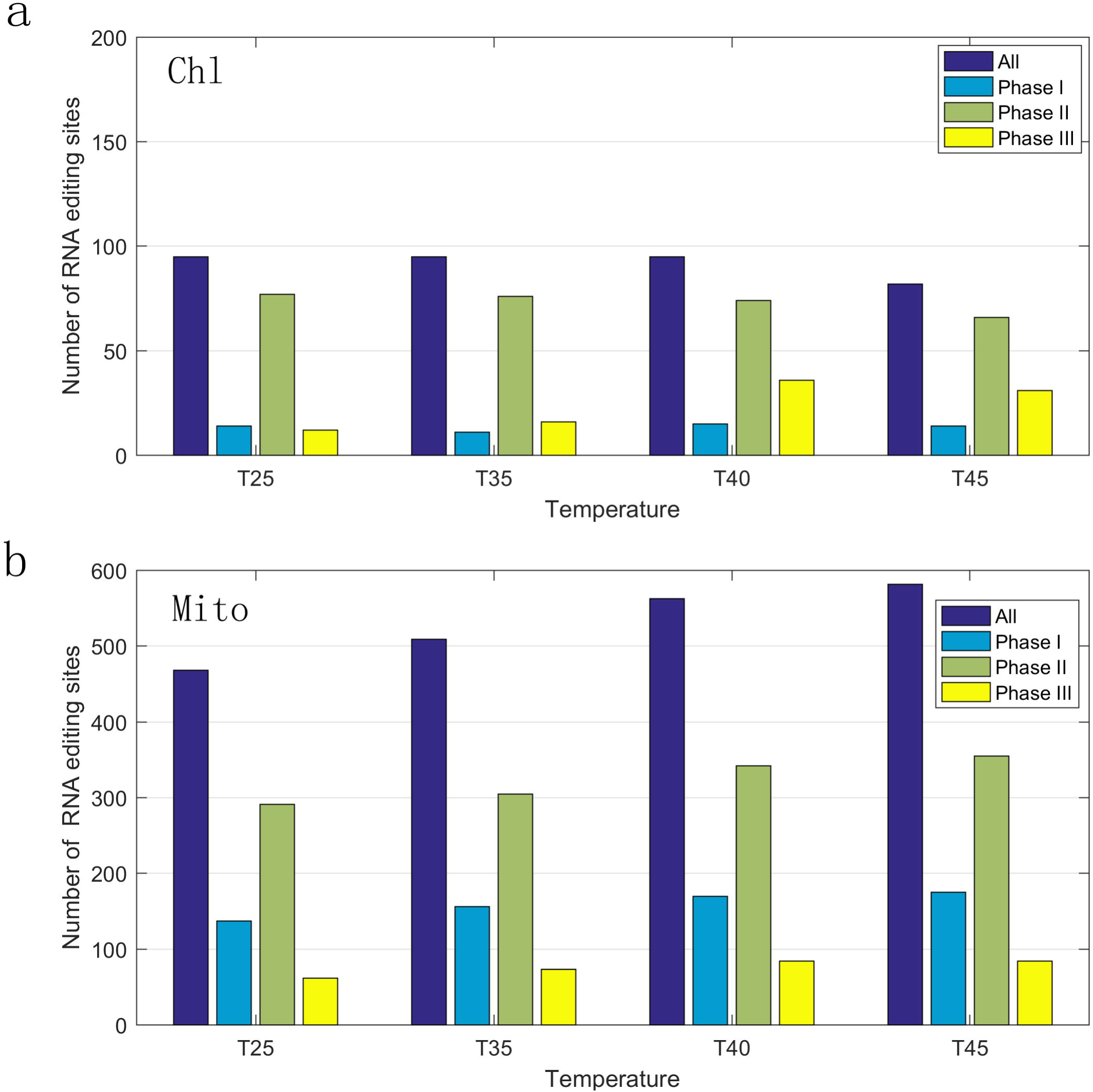
Codon position statistics of RNA editing sites. Codon position statistics of RNA editing sites were shown in a (Chl: chloroplast) and b (Mito: mitochondria) respectively.

### Reduced RNA editing efficiency with the temperature rises

We also performed statistics and cluster analysis against the RNA editing efficiency. On the whole, the average efficiency of RNA editing sites was about 0.51 in both organelles. For chloroplast and mitochondrion, the average RNA editing extent were 0.59, 0.58, 0.48, 0.42 and 0.64, 0.61, 0.58, 0.57 respectively under four conditions (25°C, 35°C, 40°C, and 45°C). With the increase of temperature, the editing extent both declined gradually, as shown in Fig.3. The results indicated that the increased temperature affected the RNA editing efficiency significantly. In addition, we searched the RNA editing sites with step-up and step-down editing extent, a total of 245 sites editing sites were identified in both organelles, and most sites have step-down editing extent. There were 175 sites demonstrated the trend of decreasing step by step (30 for chloroplast, 145 for mitochondrion), whereas 71 sites demonstrated the trend of increasing (11 for chloroplast, 60 for mitochondrion). Furthermore, the cluster analysis results also showed that the clustering relationship among samples agreed with the changes of temperatures, and there is a quiet of area demonstrated the trend of decreasing, as shown in Fig.4. Our results suggest that RNA editing is acutely sensitive to temperature, and that this response is partially affected by the thermo-sensitive secondary and tertiary RNA structures that direct editing. It is possible that differential RNA editing is one process that allows plants such as grape to rapidly adapt to varying environmental temperatures. Detailed information of RNA editing efficiency were listed in Table S2 and S5. Pairwise comparison of editing allele proportion in all samples were listed in Table S3 and S6.

**Fig.3.**
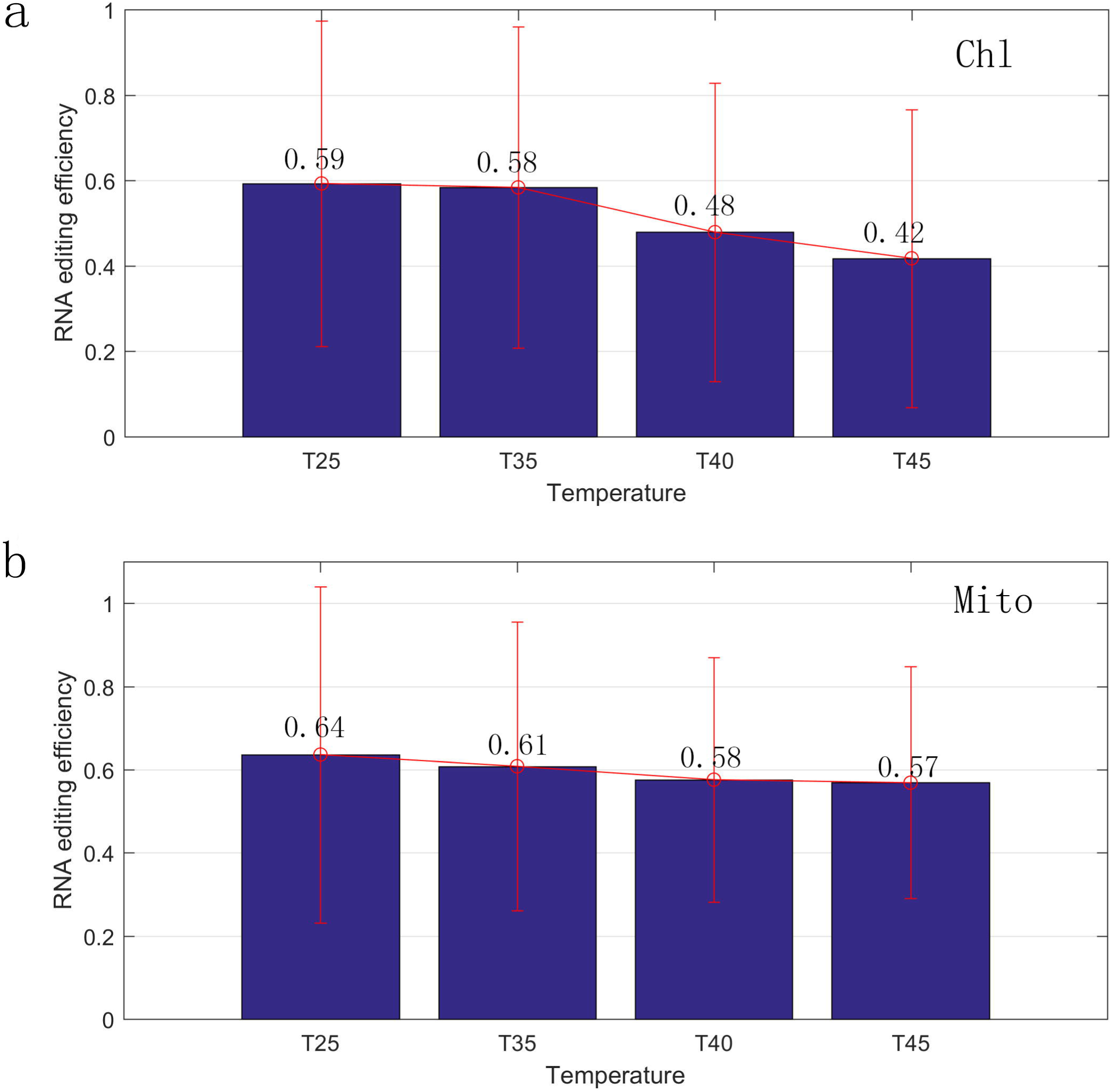
Reduced RNA editing efficiency with the temperature rises. RNA editing efficiency were shown in a (Chl: chloroplast) and b (Mito: mitochondria) respectively.

**Fig.4.**
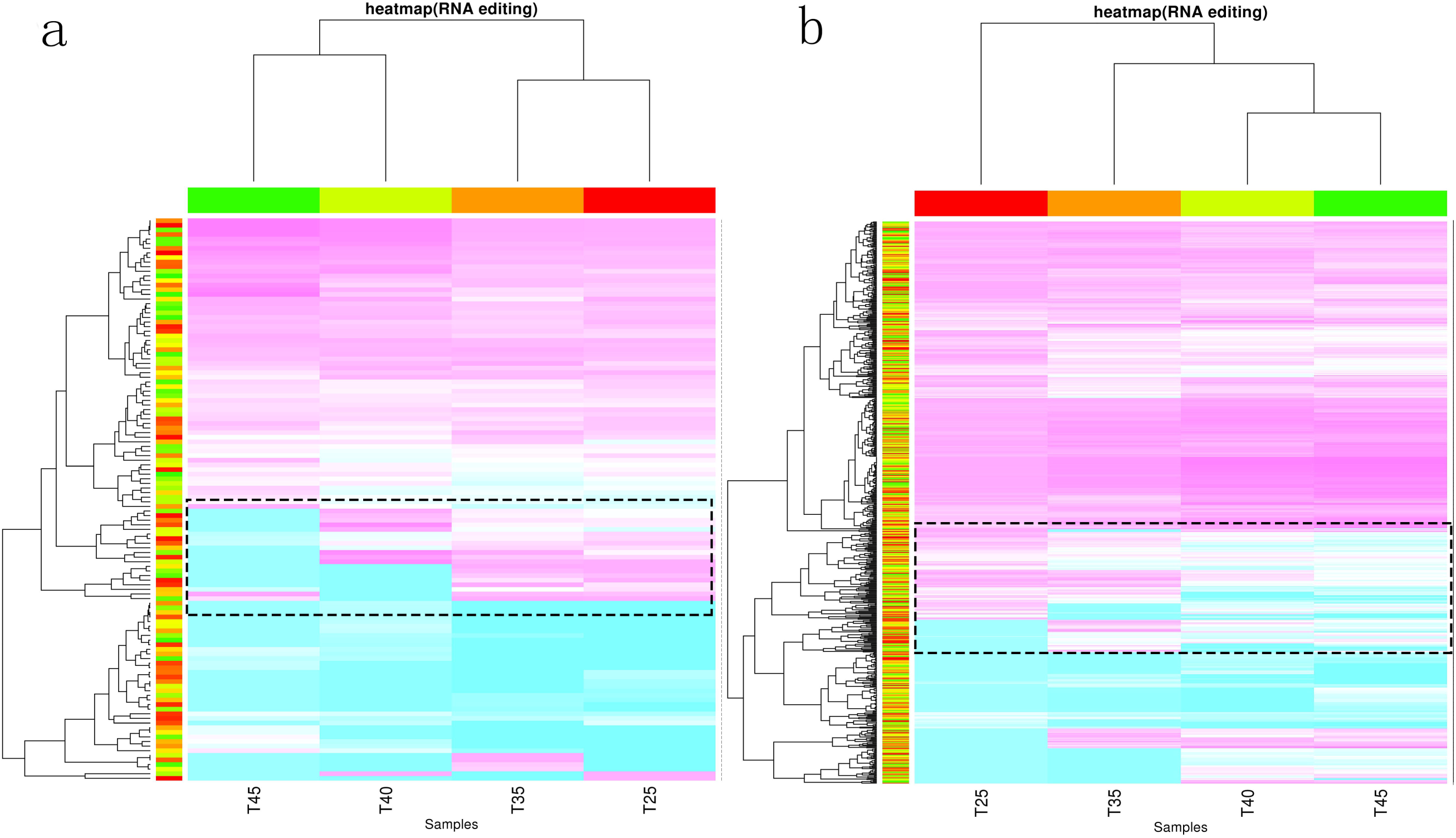
Hierarchical cluster analysis of RNA editing sites under four temperature conditions (25°C, 35°C, 40°C, and 45°C) according to the RNA editing proportions. RNA editing efficiency profiles were shown in a (Chl: chloroplast) and b (Mito: mitochondria) respectively. The x axis represents different samples, and the y axis represents editing sites. Editing sites with reduced efficiency are indicated by red dotted box, as shown in Fig 5 and Fig S1.

### Genes with changes of RNA editing efficiency

We annotated the involved genes with editing sites of step-up and step-down RNA editing extent Hence, a total of 68 genes were annotated (25 for chloroplast, 43 for mitochondrion), as shown in Fig.5 and Fig.S2. For chloroplast, several genes has more editing sites, especially *maturase K* (*matK*) and *NADH dehydrogenase subunit 2* (*ndhB*) genes, both genes have four changed editing sites. All the sites of *ndhB* gene (Chl-100212, Chl-148651, Chl-101400, Chl-101409) were C-to-U editing type, and demonstrated the trend of decreasing, the corresponding amino acid changes were His-to-Tyr, Pro-to-Leu, Ser-to-Phe, Pro-to-Leu, the four amino acids all changed to be hydrophobic. Whereas, three sites of *matK* gene demonstrated the trend of decreasing, one site showed rising trend. For mitochondrion, more genes have editing efficiency changed sites, such as *nad* gene family (*nad4/5/7*), *atp* gene family (*atp6/9*), *ccmB* gene family (*ccmB/C/FC/FC/FN*), *cox* gene family (*cox1/2/3*) and *rps* gene family (*rps4/7*).

**Fig.5.**
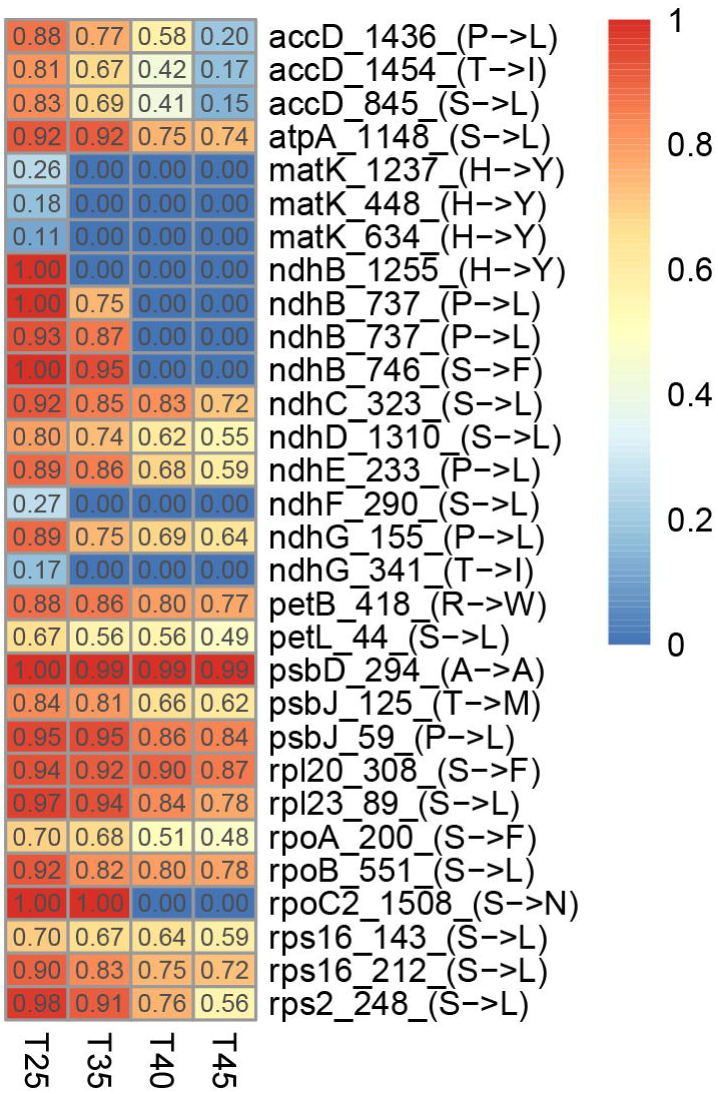
Reduced efficiency patterns of RNA editing in chloroplast. The number of RNA editing efficiency were indicated inside the box, the name of editing sites is concatenated by gene symbol, site of position and type of amino acid change.

*ndhB* gene, encodes components of the thylakoid *ndh* complex which purportedly acts as an electron feeding valve to adjust the redox level of the cyclic photosynthetic electron transporters. *ndhB* gene contains by far the higher number of editing sites, 10 in Arabidopsis, probably because the proof-reading mechanism that ensures identical sequences of the two inverted repeated regions of plastid DNA makes improbable the fixation of C-to-U back mutations. Previous studies reported that *ndh* complex is related to stress resistance, transgenic tobaccos defective in the *ndhB* gene have impaired photosynthetic activity at actual but not at high atmospheric concentrations of CO2 (Horvath et al. 2000). Furthermore, positive selection was detected in ferns and angiosperms, the adaptive evolution may affect the energy transformation and light-resistant, notably, many *nah* genes were lost or pseudogenes in gymnosperm.

*matK* gene, single copy with the length of 1,500bp, usually encodes in the *trnK tRNA* gene intron, probably assists in splicing its own and other chloroplast group II introns (Hao da et al. 2010), involving genes include the transcripts of *trnK*, *trnA*, *trnI*, *rps12*, *rpl2* and *atpF* and so on, tRNAs and proteins produced by these genes are essential for chloroplasts to function properly. Similarly, *matK* gene also suffers adaptive evolution in angiosperms, which means a lot for the transcription process of related genes (Hao da et al. 2010). Thus the changes of amino acid sites resulting from evolution or RNA editing may fine-tunes maturase performance.

### Expression analysis of RNA editing genes and PPR genes

PPR proteins are protein family that is exclusively expanded in plants, with over 450 members in *Arabidosis thaliana* and *Oryza sativa* (Lurin et al. 2004).Virtually all the investigated PPR proteins are located in either plastids or mitochondria. It is generally acknowledged that PPR proteins function as RNA-binding proteins that specially bind to the cis element of the target RNA. PPR proteins have been shown to play various roles in organelle gene expression, including transcription, RNA splicing, RNA editing, RNA cleavage, RNA stabilization and translation (Barkan and Small 2014). PPR mutants display various developmental defects (Saha et al. 2007). In order to investigate the reason for the reduced RNA editing efficiency with the increasing of temperature, we evaluated the expression of RNA editing genes and PPR proteins. However, There was no difference in the expression under different temperatures for RNA editing genes. Whereas, interestingly, a total of 8 PPR genes were found expressed in all the samples and their expression level declined significantly with the increasing of temperature. Hence, there is a positive correlation between the PPR genes expression and RNA editing efficiency, then it’s reasoned that the reduced RNA editing efficiency may result from the dropped expression of PPR genes.

**Fig.6.**
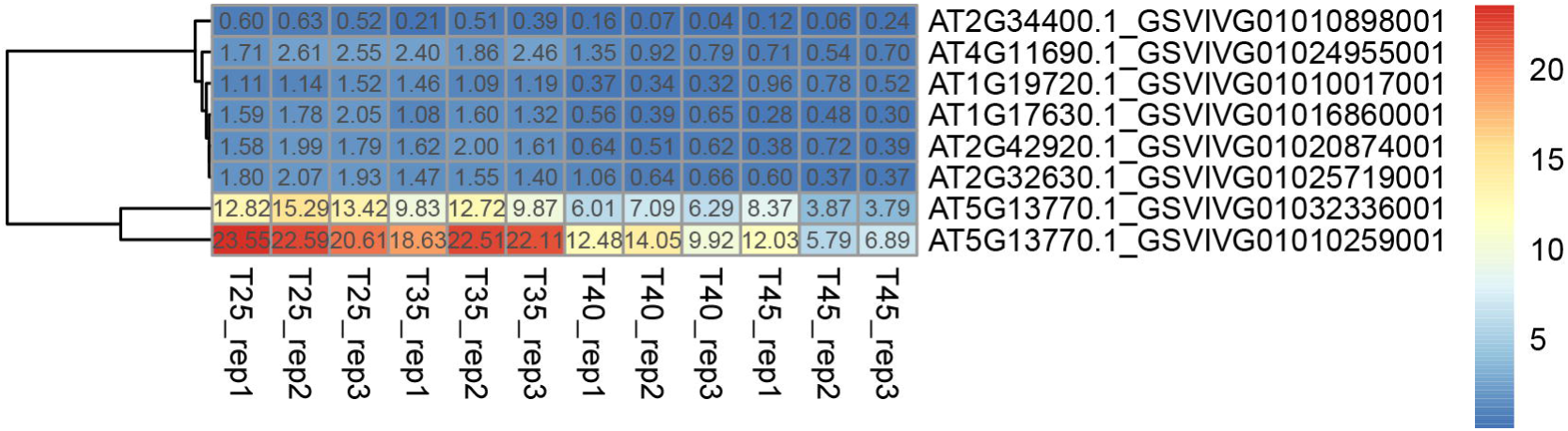
Expression patterns of PPR protein under four temperature conditions with three replicates.

## Discussion

RNA editing is an important epigenetic mechanism by which genome-encoded transcripts are modified by substitutions, insertions and/or deletions, it diversifies gnomically encoded information to expand the complexity of the transcriptome. It was first discovered in *kinetoplastid protozoa* followed by its reporting in a wide range of organisms (Maslov et al. 1994). With the advent of sequencing technology, RNA editing sites were identified in more and more organisms based on RNA deep-sequencing, especially in plants. In our study, we characterized hundreds of RNA editing sites, and the statics of editing types indicated that RNA editing typically occurs as C-to-U conversion in translated regions of organelle (mitochondrial and chloroplast) mRNAs. Most of the C-to-U changes in the protein coding regions tend to locate at first, second positions, and the physicochemical property of amino acids were mostly modified. In addition, consistent with previous studies, we also found that amino acids changes tend to be hydrophobic, therefore, plant RNA editing is believed to act as an additional proofreading mechanism to generate fully functional proteins (Ichinose and Sugita 2017; Simpson and Maslov 1999; Takenaka et al. 2013).

RNA editing has various biological functions, it can promote RNA splicing by affecting the intron structures (Castandet et al. 2010; Farre et al. 2012). Some of those editing events regulate RNA degradation and microRNA (miRNA) function. In plant, RNA editing of gene transcripts also plays a central role during plant development and evolutionary adaptation. The alteration of editing at a specific site of a mitochondrial gene can harmfully impact plant growth, development, fertility, and seed development (Kim et al. 2009; Liu et al. 2013; Sun et al. 2015; Toda et al. 2012; Yap et al. 2015). Some evidences also suggested that environmental factors, e.g., rice to the cold and maize to the heat (Kurihara-Yonemoto and Kubo 2010; Nakajima and Mulligan 2001) affect RNA editing. In addition, RNA editing has its significance during evolution (Fujii and Small 2011) and has been suggested to play a role in plant adaptation to land conditions (e.g., extreme temperatures, UV, and oxidative stress) when plants colonized the land (Fujii and Small 2011; Hammani and Giege 2014). In our study, the response of reduced RNA editing efficiency to high temperature also confirmed the relationship between environmental factors and RNA editing. It’s reasonable that RNA editing may play roles in response to environmental stress through changing the corresponding gene functions. Our results also indicated that RNA editing was more prevalent at lower temperatures, which is also accord with a previous study in animal, that the phenotypic consequences of *ADAR* (RNA editing factors in animal) deficiency in *Drosophila melanogaster* indicated that RNA editing plays an integral role in temperature adaptation by sensing and acting globally on RNA secondary structure (Buchumenski et al. 2017). It is possible that differential RNA editing is one process that allows poikilothermic animals and higher plants, such as fly and grape, to rapidly adapt to varying environmental temperatures.

Natural DNAs are usually limited to double-stranded helical shapes, whereas RNA is different, the repertoire of possible RNA secondary and tertiary structures appears limitless. RNA secondary structure is strongly correlated with function, For an RNA molecule, its structure and corresponding thermodynamic stability both contribute to functional regulation (Bonetti and Carninci 2012). Dynamic RNA structures are acutely responsive and fundamentally sensitive to abiotic factors, such as temperature and metal ion concentration, hence, it is this mutability of RNA structure that allows RNA to act as a sensor and elicit rapid cellular responses (Wan et al. 2011). Our results suggest that RNA editing is acutely sensitive to temperature, and that this response is partially affected by the thermo-sensitive secondary and tertiary RNA structures that direct editing. However, the molecular determinants underlying temperature-dependent RNA editing responses still need further study.

A number of factors are involved in plant RNA editing especially PPR and MORF proteins. Since the first PPR protein that encoded in the nuclear genome was identified and functionally described, PPR proteins have been proven to play a central role in the process of RNA editing (Hammani and Giege 2014). In our study, we show that the expression of PPRs is dramatically decreased at elevated temperatures, partially, but not fully, explaining some RNA editing sites responses to temperature.

## Conclusion

C-to-U RNA editing is a highly conserved process that post-transcriptionally modifies mRNA, generating proteomic diversity. However, its potential role in response to different stressors (heat, salt and so on) and growth development remains unclear. Our study suggest that RNA editing is responsive to environmental inputs in the form of temperature alterations. Using the angiosperms grape, we identified 123 and 628 RNA editing sites in chloroplast and mitochondria respectively with the average editing extent nearly ~60%, and detected that acute temperature alterations within a normal physiological range result in substantial changes in RNA editing levels. Additionally, The analyses also revealed that number of non-synonymous editing sites were higher than that of synonymous editing sites, and the amino acid substitution type tend to be hydrophobic. The response of reduced RNA editing efficiency to temperature alterations further confirmed the relationship between environmental factors and RNA editing, which might be through intrinsic thermo-sensitivity of the RNA structures that direct editing, or due to temperature sensitive expression of the RNA editing enzyme (*PPR* genes). Environmental cues, in this case temperature, rapidly reprogram the grape organelles transcriptome through RNA editing, presumably resulting in altered structure or function of edited proteins. However, the underlying molecular mechanisms of stress-adaptation for RNA editing still require further investigation.

## Availability of supporting data

Supporting data are included as additional files.

## Additional files

**Additional file 1: Table S1.** Information of RNA editing sites in all chloroplast samples.(XLSX)

**Additional file 2: Table S2.** A matrix for editing allele proportion of RNA editing sites in all chloroplast samples. (XLSX)

**Additional file 3: Table S3.** Pairwise comparison of editing allele proportion in all chloroplast samples. (XLSX)

**Additional file 4: Table S4.** Information of RNA editing sites in all mitochondria samples.(XLSX)

**Additional file 5: Table S5.** A matrix for editing allele proportion of RNA editing sites in all mitochondria samples. (XLSX)

**Additional file 6: Table S6.** Pairwise comparison of editing allele proportion in all mitochondria samples. (XLSX)

**Additional file 6: Fig.S1.** The attributes of RNA editing sites in chloroplast illustrated by samples at 25°C temperatures. (DOCX)

**Additional file7: Fig.S2.** Reduced efficiency patterns of RNA editing sites in mitochondria. (DOCX)

## Authors’ contributions

ADZ, XHJ, FPZ and XJZ conceived and designed the experiments, ADZ performed data analysis, ADZ, and XJZ wrote the manuscript. XHJ, TFW and XJZ provided many critical suggestions. All authors reviewed the manuscript.

## Acknowledgments

This work was funded by the National Natural Science Foundation of China (No. 61402457, No. 31702322), CAS Pioneer Hundred Talents Program and the National Project of Cause Control Theory (No.1716315XJ00200303). This work was also supported by the National Defense Science & Technology Innovation Zone Project.

## Abbreviations

A-to-I: Adenosine-to-inosine
C-to-U: Cytosine-to-uracil
ADAR: Adenosine deaminases acting on RNA
PPR: Pentatrico peptide repeat
matK: maturase K
ndhB: NADH dehydrogenase subunit 2
MORF: multiple organelle RNA editing factors

