## supplementary for "Dynamic response of RNA editing to temperature in Grape by RNA deep-sequencing"

Wuhan Botanical Garden,

Chinese Academy of Sciences,

Wuhan 430072, China

**Figures:**

**Fig.S1** The attributes of RNA editing sites in chloroplast illustrated by samples at 25°C temperatures. The density of alternative allele proportion and sequencing error proportion, the density of total read depth and average read depth, the density of read proportion for alternative allele and reference allele, the density of distance to adjacent candidate editing site, the density of log10 LLR, the density of p value, the density of GC content, the boxplot of alternative allele proportion, the boxplot of sequencing error proportion, the boxplot of total read depth, the boxplot of total read depth/average read depth, the boxplot of distance to adjacent candidate editing site, the boxplot of log10 LLR, the boxplot of p value and the boxplot of GC content are shown for RNA editing sites; GC, GC content; LLR, likelihood ratio.

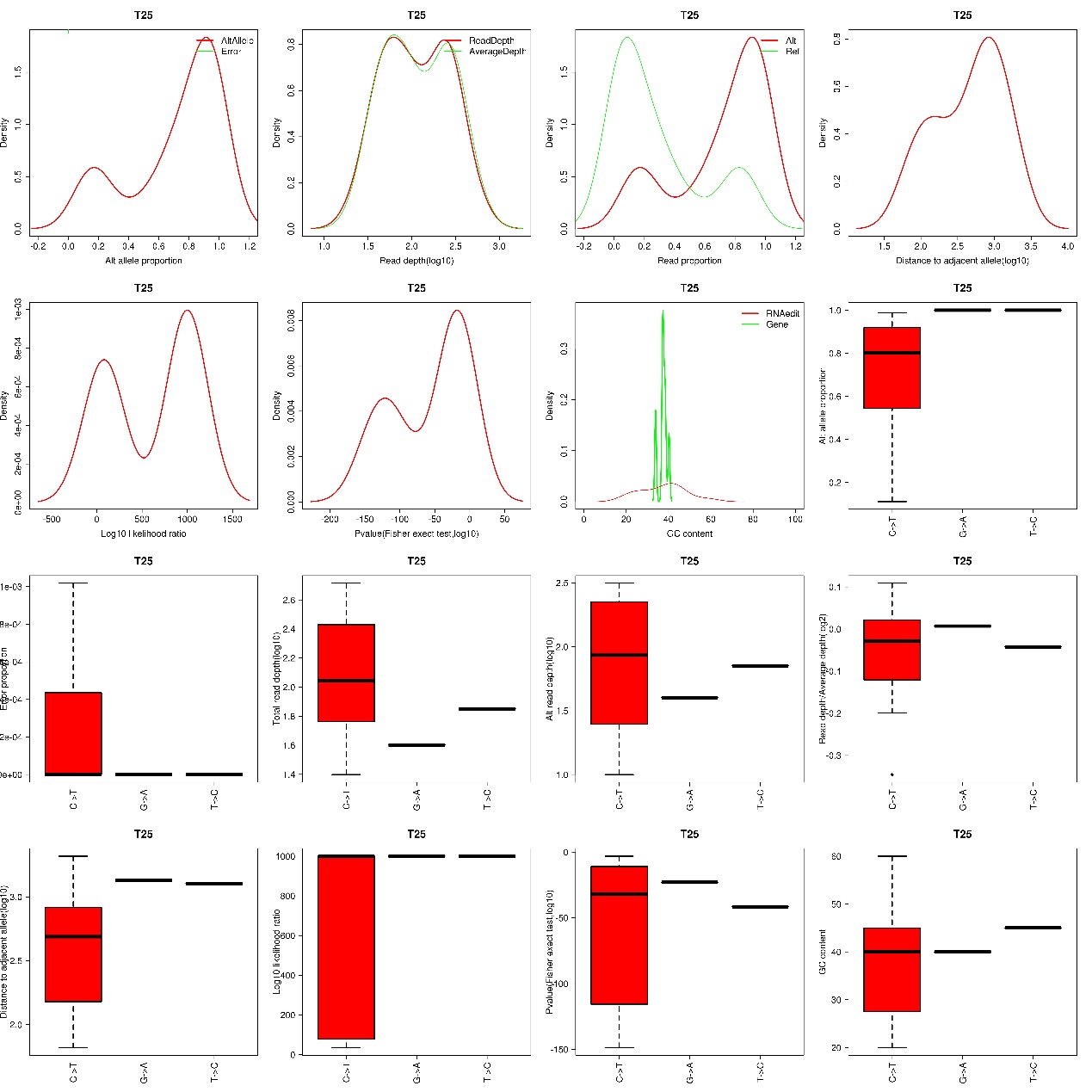

**Fig.S2** Reduced efficiency patterns of RNA editing sites in mitochondria. The number of RNA editing efficiency were indicated inside the box. The name of editing sites is concatenated by gene symbol, site of position and type of amino acid change.

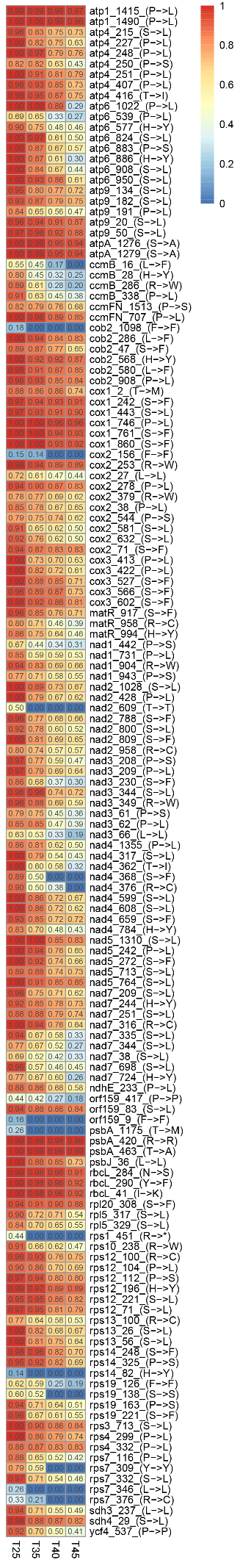
